# Treatment with 2-phospho-L-ascorbic acid mitigates biochemical phenotypes of heme oxygenase 1 deficiency

**DOI:** 10.1101/2024.07.05.602073

**Authors:** Lea-Sophie Berendes, Petra Schulze Westhoff, Ann-Marie Tobinski, Victoria Wingert, Saskia Biskup, Anja Seelhöfer, Veerle Van Marck, Barbara Heitplatz, Helmut Wittkowski, Anibh M. Das, Luciana Hannibal, Witold N. Nowak, Alicja Józkowicz, Luisa Klotz, Georg Varga, Thorsten Marquardt, Julien H. Park

## Abstract

Heme oxygenase 1 (HO-1) deficiency is a fatal genetic disorder characterized by impaired heme catabolism, leading to excessive oxidative damage and cell death. Despite evidence from non-human models suggesting mitochondrial dysfunction, the precise pathomechanisms in humans remain unclear, resulting in a lack of effective treatments. Using patient-derived lymphoblastoid cells and HO-1 knockout HEK293T cell models, we demonstrate that HO-1 deficiency is associated with altered mitochondrial morphology and impaired mitochondrial function. Furthermore, it is linked to significant ascorbic acid depletion, accompanied by compensatory upregulation of SVCT2, a key ascorbic acid transporter. Treatment with 2-phospho-L-ascorbic acid, a stable vitamin C analog, restores intracellular ascorbic acid levels and protects cells from hemin-induced cytotoxicity, highlighting its potential as a novel therapeutic strategy for HO-1 deficiency. Our study underscores the critical role of oxidative stress and mitochondrial dysfunction in HO-1 deficiency, paving the way for targeted interventions in this devastating disorder.

## INTRODUCTION

Protein-bound heme plays vital roles in physiology that include oxygen transport by hemoglobin, electron transfer in the mitochondrial respiratory chain, and nitric oxide signaling by guanylate cyclase (1, 2). In contrast, heme in its free state is a pro-oxidant molecule that can lead to cytotoxicity. Consequently, free heme is broken down by a highly efficient catabolic pathway initiated by one of two isoforms of heme oxygenase (inducible HO-1 and constitutive HO-2) (3, 4), which convert heme into equimolar amounts of carbon monoxide (CO), ferrous iron (Fe2+), and biliverdin (5). The products of this reaction themselves can exert antioxidant effects (6), while HO-1 activation stimulates various oxidant-responsive transcription factors (7). These properties make heme oxygenases key players in the response to redox stress.

The absence of HO-1 function results in an altered and progressive phenotype in murine models, highlighting the physiological importance of this protein. These animals exhibit impaired fertility, increased prenatal lethality and reduced stress resistance, accompanied by anemia, inflammation, iron accumulation, and heightened oxidative damage (8, 9). Human HO-1 deficiency (OMIM #614034), first described by Yachie and colleagues (10), mirrors this murine phenotype but is much more severe, with hemolytic anemia, chronic inflammation, and recurrent, treatment-resistant episodes of hyperinflammation. These inflammatory episodes cause extensive damage across multiple organ systems, leading to death during childhood (11). It is worth noting that although abnormal iron metabolism was common to human and murine HO-1 deficiency, dysfunction of endothelium and monocyte seemed to be far more significant in humans (11, 12).

While macrophage repopulation or bone marrow transplantation have been proposed as potential treatments, the lack of effective, clinically available therapies for HO-1 deficiency remains a significant challenge (9, 13). This treatment inadequacy is partly due to a lack of mechanistic insight into the condition’s pathophysiology. Although oxidative damage is believed to play a central role (11), the specific molecules and metabolites involved are not well understood. Studies in non-human models have implicated altered mitochondrial function as an important downstream consequence of HO-1 deficiency (14–18). However, the relevance of these changes in human HO-1 deficiency are still unclear. The initial stable period frequently observed in human cases of HO-1 deficiency suggests the presence of compensatory mechanisms that could be targeted for new therapeutic approaches.

Here, we investigated metabolic changes and compensatory mechanisms in a patient-derived lymphoblastoid cell line and HEK293T knockout cells. Our findings indicate that HO-1 deficiency is associated with ascorbate depletion, leading to increased oxidative damage. HO-1 deficient cells exhibited altered mitochondrial ultrastructure and impaired mitochondrial function. In hemin-stimulated HO-1 deficient cells, 2-phospho-L-ascorbic acid (AA2P) normalized cell viability, an effect inhibited by the ascorbate uptake inhibitor sulfinpyrazone. In conclusion, we propose ascorbic acid as a potential therapeutic agent for HO-1 deficiency.

## RESULTS

### Patient-derived LCL and HEK293T knock-out cells recapitulate the phenotype of HO-1 deficiency

To investigate the disease mechanisms associated with HO-1 deficiency in both patient-specific and genetically engineered cellular models, we utilized patient-derived lymphoblastoid cells harboring two truncating HO-1 variants resulting in a loss of protein [c.55dupG (p.Glu19Glyfs*14); c.262_268delinsCC (p.Ala88Profs*51)] (19) (LCL ^HO-1^ ^MUT/MUT^) and a wild-type control line (LCL ^HO-1^ ^WT/WT^), along with HEK293T cells genetically modified to lack HO-1 (HEK293 ^HO-1-/-^) and their wild-type equivalents (HEK293 ^HO-1^ ^+/+^), respectively. These cells were exposed to varying concentrations of hemin for 24 hours at 37 °C in a 5% CO_2_ atmosphere. The LCL ^HO-1^ ^MUT/MUT^ cells demonstrated significantly reduced viability at hemin concentrations of 50, 75, and 100 µM, as evidenced by flow cytometry, compared to cells from healthy individuals (Fig. 1A). Similarly, the HEK293 ^HO-1-/-^ cells showed markedly lower viability than their wild-type counterparts at hemin concentrations of 30 and 60 µM (Fig. 1B). These results validate the suitability of both cell models for further detailed investigation.

**Figure 1 -.**
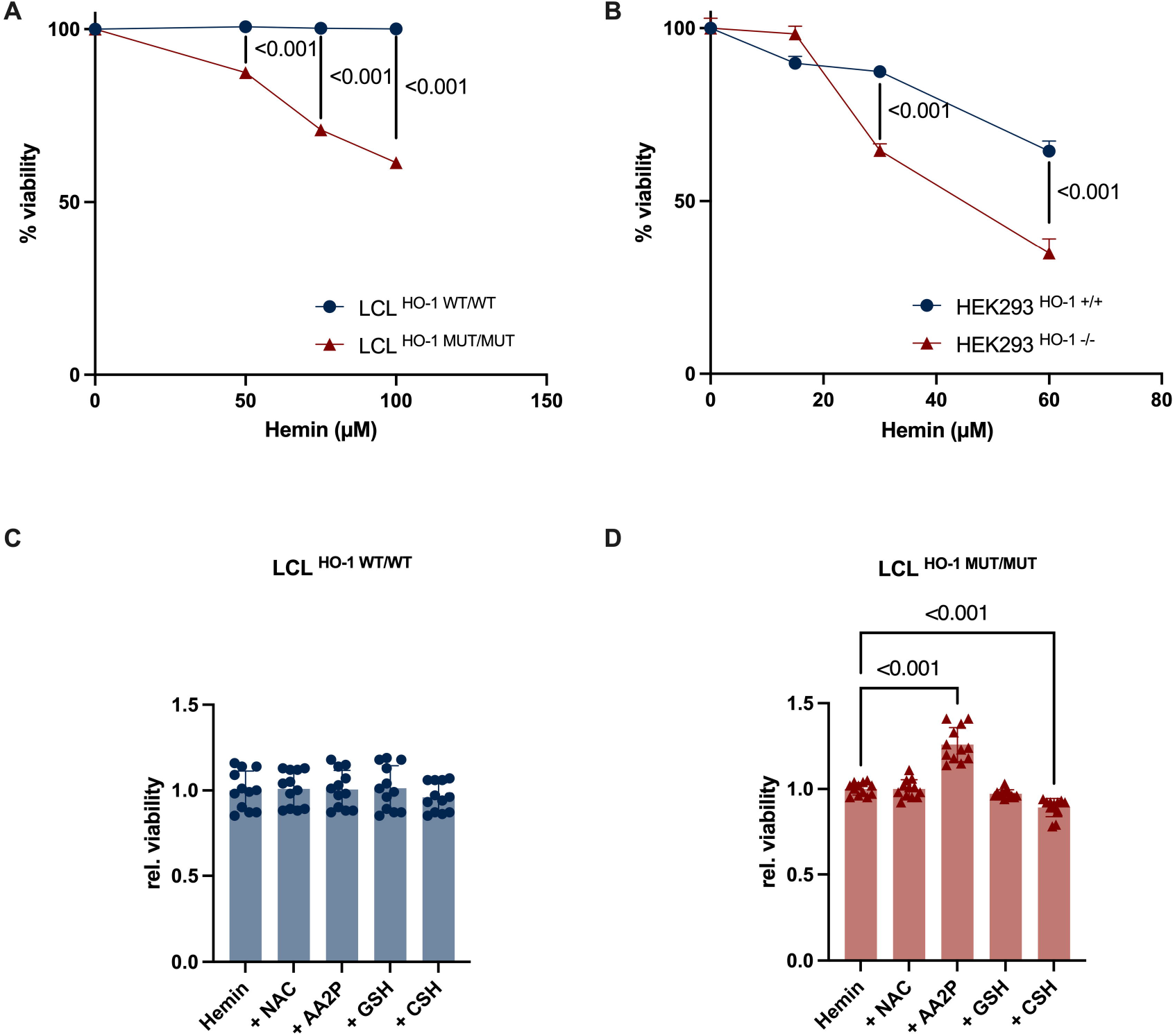
Cellular models of HO-1 deficiency replicate heme induced toxicity that is mitigated by 2-phospho-L-ascorbic acid. (A) Patient-derived LCL ^HO-1^ ^MUT/MUT^ and wild-type controls were incubated for 24 h with 0, 50, 75, and 100 µM hemin followed by viability analysis using annexin V and 7-AAD staining and subsequent flow cytometry. For normalization of data, values were divided by the mean viability of wild-type cells. (B) HEK293 ^HO-1+/+^ wild-type and HEK293 ^HO-1-/-^ cells were incubated with 0, 15, 30, and 60 µM hemin followed by identical downstream processing. (C-D) LCL ^HO-1^ ^WT/WT^ and LCL ^HO-1^ ^MUT/MUT^ were preincubated with various antioxidants (NAC - N-acetylcysteine, AA2P - 2-phospho-L-ascorbic acid, GSH - reduced glutathione, CSH - cysteamine) for 1 h followed by incubation with 75 µM hemin for 24 h. Cell viability was determined by annexin V and 7-AAD staining. Data were normalized to the viability of cells incubated with solute alone. Data are mean ± s.e.m and means were compared by ordinary two way ANOVA followed by Šídák’s post-hoc test (A-B) and ordinary one way ANOVA followed by Dunnett’s post-hoc test (C-D). Experiments were performed in *n* = 3 biological replicates with 3 technical replicates in each experiment.

### SVCT2 is upregulated in HO-1 deficient LCL, while ascorbic acid levels are decreased

To elucidate the gene expression changes associated with HO-1 deficiency, we conducted a comparative analysis using RNA sequencing (RNA-seq) between patient-derived LCL ^HO-1^ ^MUT/MUT^ cells and their wild-type equivalents. Among differentially expressed genes, this analysis identified *SVCT2* as a gene with higher expression in LCL ^HO-1^ ^MUT/MUT^ (Supplemental Fig. S1). Given its role as a principal ascorbate transporter, SVCT2 is a potential mitigator of oxidative damage (20). Subsequent validation at the protein level through quantitative immunoblotting revealed a significant increase in SVCT2 protein expression in LCL ^HO-1^ ^MUT/MUT^ cells compared to the LCL ^HO-1^ ^WT/WT^ (Fig. 2A, B). Despite the elevated expression of SVCT2, ascorbic acid levels in LCL ^HO-1^ ^MUT/MUT^ were substantially lower than in control cells (Fig. 2C), an observation consistent across the baseline conditions of the cell culture media (Supplemental Fig. S2). In contrast, HEK293 HO-1 ^-/-^ cells did not exhibit significant differences from controls in either SVCT2 expression (Fig. 2D, E) or ascorbic acid levels in cell lysates (Fig. 2F), though both parameters mirrored the trends observed in LCL cells.

**Figure 2 -.**
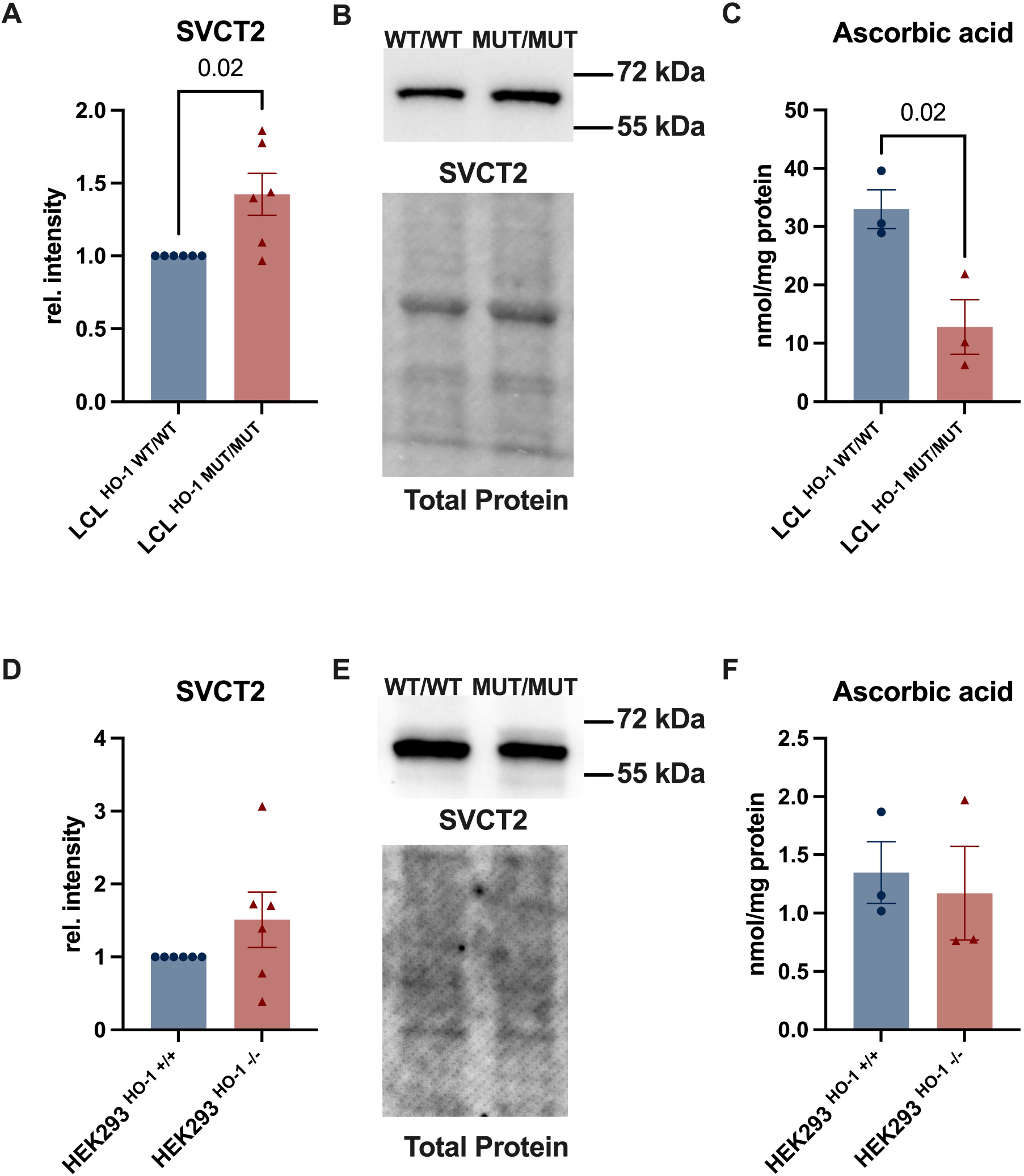
SVCT2 is upregulated in HO-1 deficient LCL while ascorbic acid levels are reduced. (A-B) LCL ^HO-1^ ^MUT/MUT^ express higher levels of the ascorbate transporter SVCT2 when compared to wild-type controls (*P*=0.02). (C) Ascorbic acid levels of cell lysates are reduced in HO-1 deficient LCL in comparison to controls (*P*=0.02). (D-E) HEK293 ^HO-1-/-^ show no statistically significant difference in SVCT2 expression in comparison to wild-type cells. (F) Ascorbic acid levels are similar between HEK293 ^HO-1+/+^ and HEK293 ^HO-1-/-^ cells. Data are mean ± s.e.m and means were compared using the two-sided nonparametric Mann-Whitney *U* test. Immunoblotting was performed in *n* = 6 replicates, while ascorbic acid measurements were performed in *n* = 3 replicates.

### HO-1 deficiency is associated with mitochondrial enlargement and altered function

Given previous, partly conflicting reports on the functional relevance of HO-1 for mitochondria (16, 21, 22), we analyzed mitochondrial structure and function in both LCL and HEK293 models of HO-1 deficiency. Transmission electron microscopy (TEM) revealed no significant differences in mitochondrial structure between patient-derived and control LCL at baseline (Fig. 3A, C). However, following stimulation with hemin (25 µM), a statistically significant enlargement of all measured parameters was observed (Fig. 3D). In HEK293 cells, HO-1 deficiency was associated with mitochondrial enlargement in all measured parameters except length at baseline, a finding that was more pronounced upon hemin stimulation (Fig. 3B, E-F).

**Figure 3 -.**
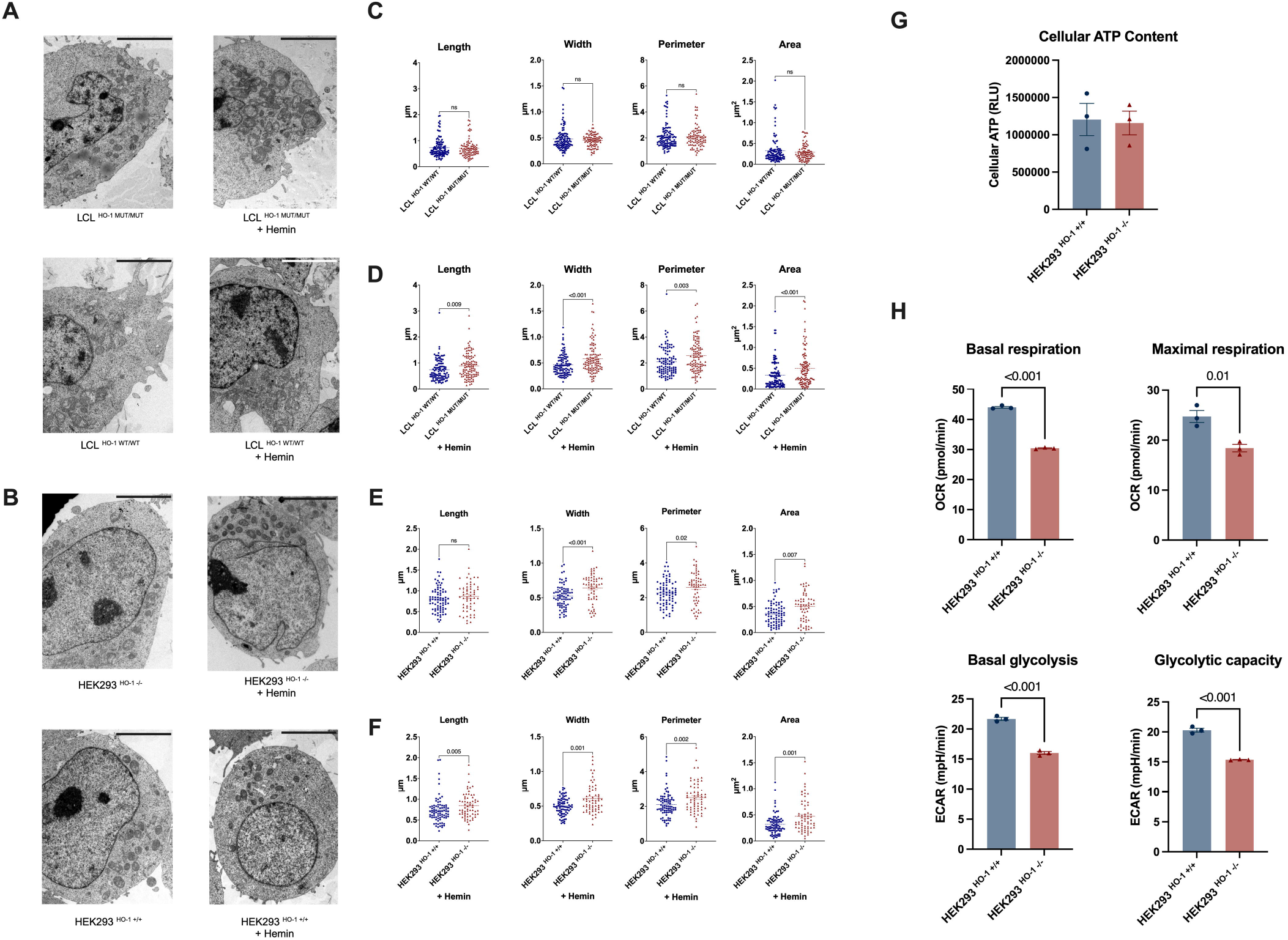
HO-1 deficiency is associated with impaired mitochondrial function and enlarged mitochondria. Transmission electron microscopy imaging of wild-type and HO-1 deficient LCL (A) and HEK293T cells (B) with and without hemin stimulation revealed no statistically significant difference of LCL ^HO-1^ ^MUT/MUT^ compared to LCL ^HO-1^ ^WT/WT^ at baseline (C). Stimulation with 25 µM hemin for 24h led to significant swelling with increased length (*P*=0.009), width (*P*< 0.001), perimeter (*P*=0.003), and area (*P*<0.001) in LCL ^HO-1^ ^MUT/MUT^. In HEK293 ^HO-1-/-^ width (*P*<0.001), perimeter (*P*=0.02), and area (*P*=0.007) were significantly increased when compared to wild-type controls at baseline (E). This swelling increased further upon stimulation with 25 µM hemin for 24h (F). Cellular ATP content as determined by luciferase assay showed no significant difference between HEK293 ^HO-1-/-^ and wild-type HEK293T cells (G). In HO-1 deficient HEK293, both basal (*P*<0.001) and maximal respiration (*P*=0.01) as determined by OCR measurements as well as basal glycolysis (*P*<0.001) and glycolytic capacity (*P*<0.001) were significantly decreased in comparison to wild-type controls (H). Data are mean ± s.e.m and means were compared using the two-sided nonparametric Mann-Whitney *U* or two-sided student’s t-test where appropriate. TEM measurements were performed in *n* = 14 replicates per cell type and condition. Mitochondrial bioenergetics analyses were performed in *n* = 3 biological replicates with 3 technical replicates in each experiment. Scale bars represent 5 µm.

On a functional level, spectrophotometrical assessment of the activity of electron transfer chain complexes in HO-1 deficient HEK293T cells revealed an increased activity of mitochondrial ATP synthase of 396 nmol/min/mg protein when compared to wild-type controls (279 nmol/min/mg protein). All other complexes as well as the activity of citrate synthase, which was used as a mitochondrial marker enzyme, were comparable to wild-type controls (Table 1). Despite this difference in ATP synthase activity, total ATP luminescence levels of HEK293 HO-1 ^-/-^ did not differ significantly from wild-type cells (*P*=0.87, Fig. 3G). HO-1 deficient cells exhibited a significantly reduced basal respiration (44.07±0.31 vs 30.43±0.17 pmol/min, *P*<0.001, Fig. 3H) and maximal respiration (24.74±1.21 vs 18.41±0.75 pmol/min, *P*=0.01). Likewise, basal glycolysis and glycolytic capacity were significantly lower in HO-1 deficient cells (Fig. 3H).

**Table 1.** Spectrophotometric analysis of respiratory chain complex activity in HO-1 deficient HEK293T cells.

|  | **HEK293**<sup>HO-1 +/+</sup><br>(nmol/min/mg protein) | **HEK293**<sup>HO-1 -/-</sup><br>(nmol/min/mg protein) |
| --- | --- | --- |
| **Complex I + III** | 43 | 54 |
| **Complex II + III** | 33 | 21 |
| **Complex IV (COX)** | 42 | 32 |
| **Complex V (ATP synthase)** | 279 | 369 |
| **Citrate synthase** | 105 | 119 |

Data are mean values of $n = 3$ technical replicates.

### 2-phospho-L-ascorbic acid (AA2P) attenuates the biochemical phenotype of HO-1 deficiency

We next explored the impact of various antioxidant compounds on cell survival in LCL and HEK293T models of HO-1 deficiency. The selection of antioxidants and their concentrations were informed by their potential for human application and findings from previous pharmacokinetic studies (Table 2) (23–26). In LCL ^HO-1^ ^WT/WT^ cells, pre-treatment with a range of antioxidants showed no significant impact on cell viability (Fig. 1C). In contrast, LCL ^HO-1^ ^MUT/MUT^ cells displayed a marked increase in viability following pre-incubation with 100 µM 2-phospho-L-ascorbic acid (AA2P), while treatment with 126 µM cysteamine resulted in reduced viability. No significant differences in viability were observed for treatments with reduced glutathione (GSH, 24 µM) or N-acetyl-cysteine (NAC, 95 µM).

**Table 2.** Antioxidant concentrations used in screening.

| Antioxidant | Concentration | Reference |
| --- | --- | --- |
| N-Acetyl-L-cysteine | 95.0 μM | Pendyala et al. Cancer Epidemiol Biomarkers Prev. 1995 Apr-May;4(3):245-51. |
| 2-Phospho-L-ascorbic acid trisodium salt | 100.3 μM | Levine et al. Proc Natl Acad Sci U S A. 1996 Apr 16;93(8):3704-9. |
| L-Glutathione reduced | 23.9 μM | Look et al. Eur J Clin Nutr. 1997 Apr;51(4):266-72. |
| Cysteamine hydrochloride | 126.4 μM | Gangoiti et al. Br J Clin Pharmacol. 2010 Sep;70(3):376-82. |

Subsequent dose-finding studies with AA2P in both LCL cell types revealed an enhancement in cell viability in the LCL ^HO-1^ ^MUT/MUT^ cells, with no noticeable effect on the WT/WT cells (Fig. 4A-C). At AA2P concentrations of 1-10 mM, no significant differences in viability were observed between the two cell types. However, a higher AA2P concentration of 20 mM led to a decline in viability in the LCL ^HO-1^ ^WT/WT^ cells, while LCL ^HO-1^ ^MUT/MUT^ maintained stable viability.

**Figure 4 -.**
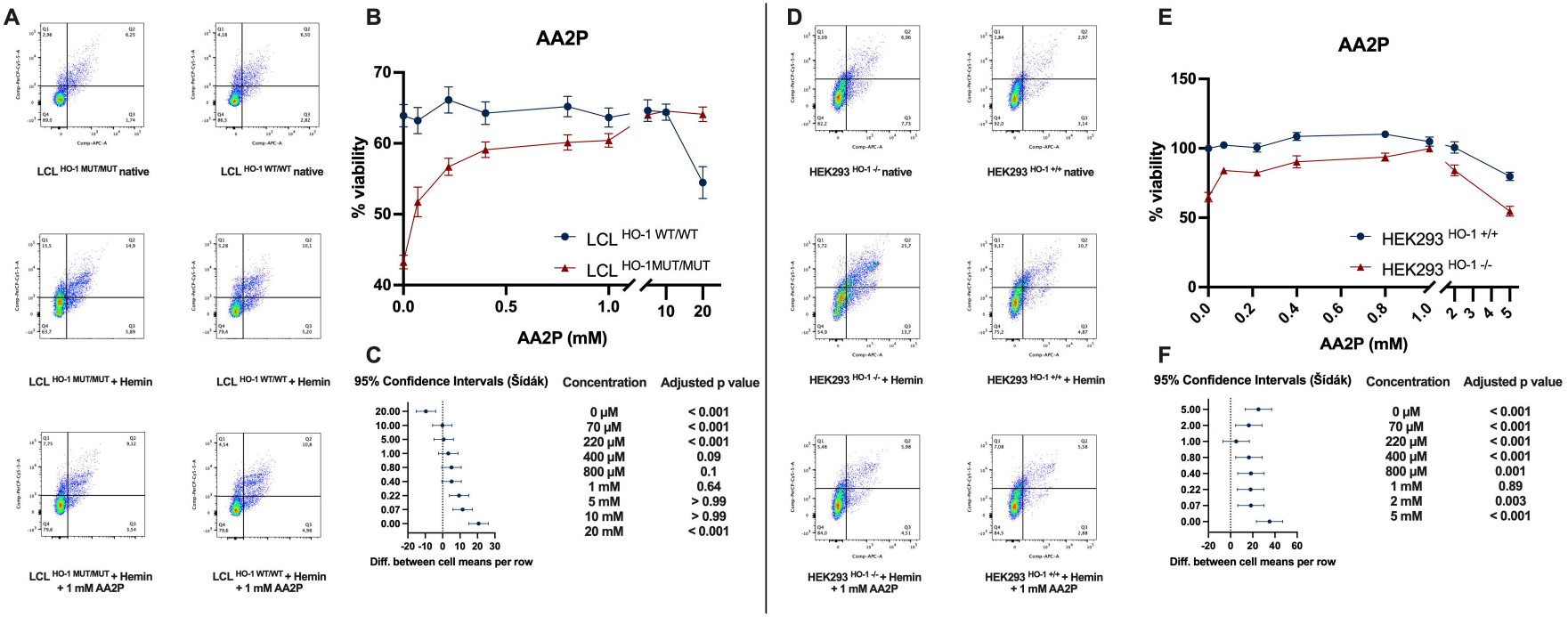
2-phospho-L-ascorbic acid improves cell survival in hemin-challenged HO-1 deficient LCL and HEK293T cells. (A-C) Treatment with ascending doses of 2-phospho-L-ascorbic acid (AA2P) attenuates hemin-induced cell death. Following pre-incubation with ascending doses of AA2P for 1 h, hemin-treated LCL showed a normalization of cell viability to levels comparable to controls. At concentrations of 400 µM, 800 µM, and 1 mM, no statistically significant difference to wild-type controls was seen (*P*=0.09, *P*=0.1, and *P*=0.64, respectively). Higher doses led to a decrease in the viability of wild-type cells, while HO-1 deficient cells were unaffected. (D-F) HO-1 deficient HEK293T cells showed improved survival upon AA2P treatment with no statistically significant difference to wild-type controls at an AA2P concentration of 1 mM (*P*=0.89). Higher doses led to a reduction in viability in both cell types. Data are presented as mean ± s.e.m. Means were compared using ordinary two-way ANOVA followed by Šídák’s post-hoc test. Experiments were performed in *n* = 3 biological replicates with 3 technical replicates in each experiment.

To determine if these effects were specific to LCL, similar dose-finding studies were conducted in the HEK293T model. In HEK293 ^HO-1-/-^ cells, AA2P preincubation improved cell viability with no statistically discernible difference at a concentration of 1 mM, aligning with observations in LCL (Fig. 4D-F). However, at higher concentrations, a reduction in cell viability was noted in both HEK293T genotypes, with the HO-1 deficient cells exhibiting lower viabilities at 10 and 20 mM AA2P respectively.

### Treatment with AA2P increases intracellular ascorbic acid concentrations regardless of genotype

To assess whether treatment with AA2P led to increased intracellular ascorbic acid levels, we compared ascorbic acid concentrations in purified cell lysates from untreated LCLs and HEK293T cells, and from cells exposed to hemin, as well as cells pre-treated with AA2P before hemin exposure (Fig. 5A, B and E, F). In LCL ^HO-1^ ^MUT/MUT^ cells, treatment with hemin led to a statistically significant reduction in ascorbic acid when compared to untreated cells (Fig. 5A). In contrast LCL ^HO-1^ ^WT/WT^ exhibited a reduction that did not reach statistical significance (Fig 5 B). This reduction in ascorbic acid levels was associated with a decrease in the GSH/GSSG ratio. Preincubation with 800 µM AA2P resulted in a statistically significant increase in ascorbic acid both in LCL ^HO-1^ ^MUT/MUT^ (70.5 ± 1.0 nmol/mg protein, *P*<0.001) and LCL ^HO-1^ ^WT/WT^ (97.27±21.78 nmol/mg protein, *P*=0.02).

**Figure 5 -.**
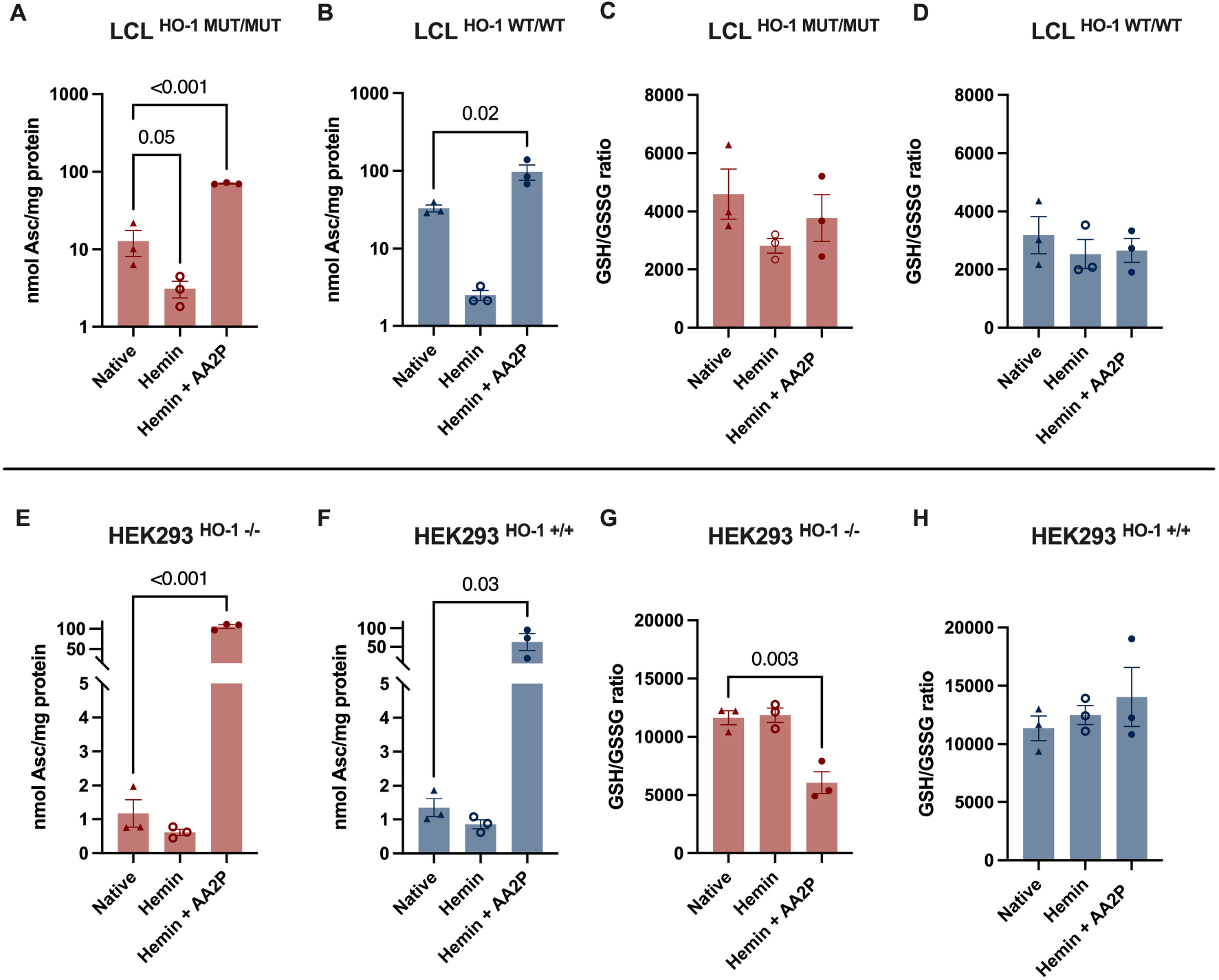
Hemin stimulation decreases intracellular ascorbate with inconsistent effects on the GSH/GSSG ratio. (A) Hemin stimulation of LCL ^HO-1^ ^MUT/MUT^ leads to a reduction in intracellular ascorbic acid (AA) (*P*=0.05). Pre-treatment with AA2P increases ascorbic acid levels above baseline (*P*<0.001). (B) While hemin stimulation decreases ascorbic acid levels without reaching statistical significance, pre-treatment with AA2P leads to a significant increase in AA levels above baseline (*P*=0.02) in LCL ^HO-1^ ^WT/WT^. (C-D) GSH/GSSG ratios show modest, non-significant reductions upon hemin stimulation in both cell lines. (E-F) No statistically significant reduction in AA concentration was observed in either HO-1 deficient or wild-type HEK293T upon hemin stimulation. Preincubation with AA2P led to a significant increase in AA levels (*P*<0.001 and *P*=0.03). (G) In HEK293 ^HO-1-/-^, the GSH/GSSG ratio remains stable upon hemin stimulation but decreases significantly following pre-treatment with AA2P (*P*=0.003). (H) No significant changes in the GSH/GSSG ratio were observed in HEK293 ^HO-1+/+^. Data are presented as mean ± s.e.m. Means were compared using ordinary one-way ANOVA followed by Dunnett’s post-hoc test. Experiments were performed in *n* = 3 replicates.

In HEK293T cells, both HEK293 ^HO-1-/-^ and HEK293 ^HO-1^ ^+/+^ exhibited about 10-fold lower levels of ascorbic acid at baseline compared to LCL. Treatment with hemin led to only a slight decline in ascorbic acid that did not reach statistical significance (Fig. 5E, F). Preincubation with AA2P effected a statistically significant increase in ascorbic acid, reaching values equivalent to those observed in AA2P-treated LCL cells.

### Cellular uptake of AA2P is required to improve cell survival in HO-1 deficiency

We next asked whether the effect of AA2P was an epiphenomenon caused by extracellular ROS scavenging rather than an intracellular effect. Therefore, we assessed the effect of AA2P on the relative viability in LCL ^HO-1^ ^WT/WT^ (Fig. 6A) and LCL ^HO-1^ ^MUT/MUT^ (Fig. 6B) cells with and without hemin stimulation in the presence of sulfinpyrazone at a concentration of 500 µM. This compound is a potent blocker of SVCT2, the principal ascorbic acid uptake transporter that is upregulated in HO-1 deficient LCL (Fig. 2 A-B). Incubation of sulfinpyrazone alone led to a decrease in relative cell viability in LCL ^HO-1^ ^WT/WT^ (0.8 vs 0.96, *P*=0.002). As expected, hemin stimulation of LCL ^HO-1^ ^MUT/MUT^ led to a significantly decreased cell viability (*P*<0.001) that was improved by pre-incubation with AA2P. Treatment with sulfinpyrazone before pre-incubation with AA2P inhibited this effect following hemin stimulation in LCL ^HO-1^ ^MUT/MUT^ (*P*<0.001). Interestingly, treatment of LCL ^HO-1^ ^WT/WT^ with sulfinpyrazone led to a significantly decreased relative viability upon AA2P and hemin incubation (0.66 vs 0.96, *P*<0.001). Therefore, inhibition of SVCT2 abrogates protective effects of AA2P on HO-1 deficient cells.

**Figure 6 -.**
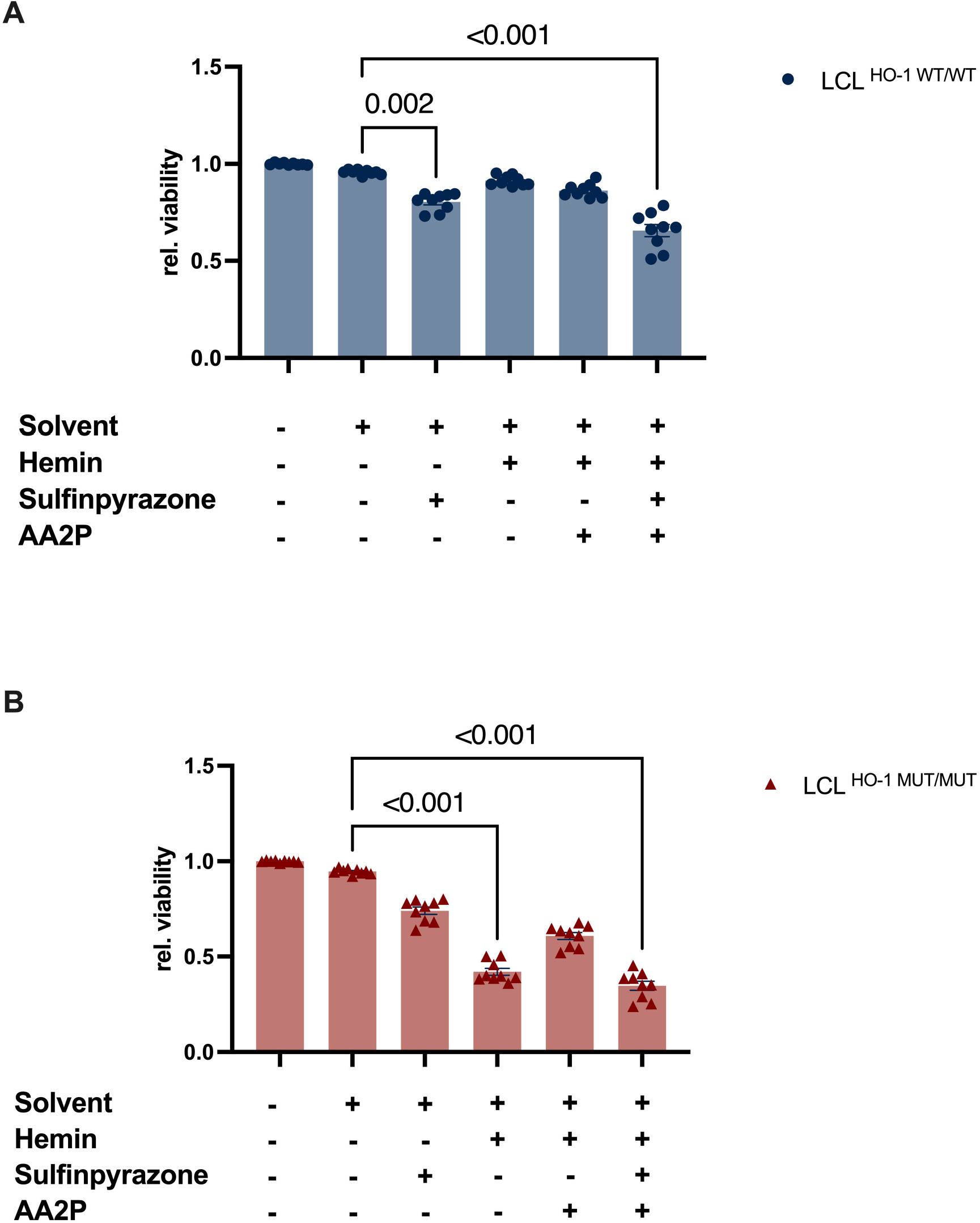
Inhibition of the ascorbic acid transporter SVCT2 counteracts protective effects of AA2P in hemin stimulated LCL. Treatment with the SVCT2 inhibitor sulfinpyrazone (500 µM) alone led to a reduction in relative viability in LCL ^HO-1^ ^WT/WT^ (*P*=0.002). Treatment with hemin led to a reduction of relative viability in LCL ^HO-1^ ^MUT/MUT^ (*P*<0.001) while no significant change was observed in LCL ^HO-1^ ^WT/WT^. Pre-incubation of LCL ^HO-1^ ^MUT/MUT^ with 1 mM AA2P increased viability as expected, an effect that was abrogated in the presence of sulfinpyrazone (*P*<0.001). In LCL ^HO-1^ ^WT/WT^, sulfinpyrazone followed by AA2P pre-incubation and hemin stimulation led to a significant reduction in relative viability (*P*<0.001). Data are presented as mean ± s.e.m. Means were compared using the Kruskal-Wallis test followed by Dunn’s post-hoc test. Experiments were performed in *n* = 3 biological replicates with 3 technical replicates in each experiment.

## DISCUSSION

Heme oxygenase 1 deficiency is a devastating condition with extremely limited treatment options and multiple failed treatment attempts with immunosuppressive drugs (11). Although hematopoietic stem-cell transplantation (HSCT) has shown promise in animal studies as a potential cure (9), current evidence from human applications is limited (13). Based on suspected pathomechanisms, *ex vivo* gene therapy might be a candidate treatment although no such attempts have been reported. This underscores the critical need for more effective treatments. A common trait among previously investigated patients is a clinically silent period with no indication of inflammatory bouts despite the presence of subclinical changes in laboratory parameters (11). This suggests the presence of currently unknown compensatory mechanisms. Here, we aimed to characterize the metabolic consequences of HO-1 deficiency and adaptive mechanisms to identify potentially therapeutic modifiers of the phenotype.

To this end, we employed both patient-derived lymphoblastoid cells (Fig. 1A) and genetically modified HEK293T cells (Fig. 1B). These models recapitulate the key features of HO-1 deficiency, facilitating detailed analysis of the associated metabolic and oxidative stress responses. Notably, LCL originate from bone marrow, a tissue particularly affected by HO-1 deficiency. The bone marrow niche plays a pivotal role in hematopoiesis by regulating hematopoietic stem cell stimulation and providing protection against stress conditions. HO-1 exerts its antioxidative and anti-inflammatory effects within the bone marrow, ensuring the functional capability of hematopoietic stem cells (HSCs) (27). Conversely, HO-1 deficiency has been demonstrated to lead to HSC exhaustion in murine models (28). The reduced viability of both LCL ^HO-1^ ^MUT/MUT^ and HEK293 ^HO-1-/-^ cells in response to hemin exposure reproduces these phenotypes, highlighting the robustness of these cellular models to study HO-1 deficiency.

We identified changes in both mitochondrial structure and function of HO-1 deficient cells. At baseline, LCL ^HO-1^ ^MUT/MUT^ showed normal mitochondrial morphology but developed enlarged mitochondria upon hemin stimulation (Fig. 3C,D), while HO-1 deficient HEK cells had larger mitochondria even at baseline, with these differences becoming more pronounced after hemin stimulation (Fig. 3E,F). These findings align with animal studies demonstrating HO-1’s role in regulating mitochondrial biogenesis and protecting against oxidative stress. In murine models, HO-1 deficiency impairs mitochondrial biogenesis and function, as evidenced by reduced mitochondrial DNA content and respiratory capacity (14–16). This study extends these findings to human cells, showing that HO-1 is essential for maintaining mitochondrial integrity, particularly under stress conditions. Functional assays further revealed that HO-1 deficient cells exhibited both significantly impaired mitochondrial respiration and glycolytic efficiency with impaired glycolysis at baseline as well as under stress. The observed upregulated ATP synthase activity (Table 1) indicates a compensatory mechanism in response to these dual impairments, underscoring the enzyme’s critical role in cellular bioenergetics (Fig. 3H). The observed enlargement of mitochondria in HO-1 deficient cells, particularly under stress, in conjunction with the apparent increase in ATP synthesis might be indicative of stress-induced mitochondrial hyperfusion (SiMH), a protective stress-response resulting in elongated mitochondria (29) as well as increased mitochondrial ATP production (30). The equal overall ATP content of HO-1 deficient HEK293T cells (Fig. 3G) might indicate successful compensation of impaired mitochondrial function by increased overall ATP synthase activity, although further research on the underlying mechanisms is warranted. Overall, these findings substantiate mitochondrial dysfunction as a major factor in the pathogenesis of human HO-1 deficiency.

With regards to compensatory mechanisms, we found a marked upregulation of SVCT2 in LCL ^HO-1^ ^MUT/MUT^, evident both at the RNA and protein level. Despite the increased expression of this principal ascorbic acid (AA) transporter (31), intracellular AA levels were markedly lower in HO-1 deficient cells compared to wild-type controls (Fig. 2C+F).The preserved AA levels in the cell culture media of these cells (Supplementary Fig. S2), suggest that the compensatory upregulation of SVCT2 upon increased oxidative stress is not sufficient to meet the elevated AA consumption, leading to a net decrease in intracellular levels. Interestingly, no significant upregulation of SVCT2 was found in HEK293 ^HO-1-/-^. This suggests cell-type specific compensatory mechanisms or susceptibility to AA oxidation, especially given the exceptionally high AA content of lymphocytes (32).

Given the upregulation of SVCT2 and decreased intracellular AA concentration of LCL ^HO-1^ ^MUT/MUT^, we evaluated the potential of 2-phospho-L-ascorbic acid (AA2P), a stable form of AA, as a therapeutic agent for HO-1 deficiency (Fig. 4). We found a significant improvement in cell survival with AA2P pre-incubation, particularly in LCL ^HO-1^ ^MUT/MUT^ cells and were able to compensate for the effects of hemin stimulation.

The concentrations of AA2P needed for this effect are considerably higher than the plasma levels of AA in healthy individuals (50-80 µM) (23, 32) but within the range found in lymphocytes (4 mM) (33). These concentrations could potentially be achieved in lymphocytes through parenteral AA administration (34).We also demonstrated that the cellular uptake of AA2P is crucial for its protective effects since inhibition of SVCT2 with sulfinpyrazone significantly reduced cell viability (Fig. 6), indicating that intracellular ascorbic acid is essential for mitigating hemin-induced toxicity.

This observation may also shed light on the reason for the differences in the severity of the HO-1 deficiency phenotype in mice and humans. Despite increased mortality in the fetal period, adult mice under control conditions do not show such significant vascular dysfunction, as humans, although the symptoms worsen with age (12).

However, mice still regularly survive for more than 1.5 years (28). Our results indicate that the alleviation of HO-1 deficiency symptoms in mice may be related to the ability to synthesize ascorbic acid, typical of mice, but lost in humans and other primates (35). Interestingly, ascorbic acid synthesis decreases with age (36), and vitamin C deficiency affects the vascular integrity of genetically modified mice unable to synthesize ascorbic acid, with potentially profound effects on the pathogenesis of vascular diseases (37).

The effect of AA2P on LCL ^HO-1^ ^MUT/MUT^ viability was accompanied by a non-significant increase in the GSH/GSSG ratio compared to cells treated with hemin (Fig. 5C), which exhibited a significant reduction compared to the baseline. This enhancement in the ratio indicates a shift towards a more reduced cellular redox state, suggesting a decrease in oxidative stress within these cells (38). In contrast, HEK293 ^HO-1-/-^ exhibit a reduced GSH/GSSG ratio following AA2P treatment (Fig. 5G). This somewhat surprising finding might, at face value, be interpreted as a pro-oxidant effect of AA2P. However, the increased survival of HEK293 ^HO-1-/-^ argues against it. Instead, it may rather reflect an increased GSH turnover, since GSH is used to regenerate AA, e. g. from DHA (39–41). The GSH/GSSG ratio, while indicative of redox changes, does not capture the entire spectrum of cellular redox processes, which involve other antioxidants and signaling pathways (42). Taking cellular compartmentalization into account, this ratio might not correctly reflect the redox state in specific intracellular areas such as the mitochondria that generate reactive oxygen species. Furthermore, inherent biological variability among cell types and conditions can influence GSH and GSSG levels (43), potentially impacting the interpretability of this ratio across different experimental setups (44). A deeper exploration is warranted to elucidate the precise mechanisms of oxidative damage linked to HO-1 deficiency. Prior research has highlighted HO-1’s role in detoxifying H_2_O_2_ (45), most likely in close relation to the enzyme’s involvement in key regulatory pathways of oxidative stress as well as hypoxia (46). Consequently, spatially resolved analysis of H_2_O_2_ concentrations and the impact of HO-1 deficiency on other redox-related genes’ regulation could yield further insights into the disorder’s pathophysiology.

In conclusion, our study identified previously unknown aspects of HO-1 deficiency, including the upregulation of SVCT2, mitochondrial alterations, and the potential therapeutic benefits of ascorbic acid. These findings advance our understanding of pathomechanisms of HO-1 deficiency and highlight new approaches for therapeutic intervention. Future studies should focus on molecular mechanisms of SVCT2 regulation and as well as the pathophysiology of mitochondrial dysfunction to mitigate the effects of HO-1 deficiency effectively. By exploring these avenues, more effective treatment strategies for HO-1 deficiency will be attained.

## METHODS

### Sex as a biological variable

Only LCL derived from a male subject were available for analysis. To account for sex- and tissue-specific effects, key experiments were replicated in HEK293T cells, which are derived from a female fetus (47, 48). The results are expected to be relevant to all humans.

### Chemicals and reagents

All chemicals were obtained from Sigma-Aldrich (St. Louis, MO, USA) unless stated otherwise. Cell culture materials were purchased from Thermo Fisher Scientific (Waltham, MA, USA).

### Generation of cellular models of HO-1 deficiency

To study the phenotype of HO-1 deficiency in detail, we generated a HO-1 deficient lymphoblastoid cell line (LCL). Clinical details as well as the extensive preparation protocol have been published elsewhere (19). In brief, peripheral venous EDTA blood samples were collected from a patient carrying two pathogenic variants in *HMOX1* (c.262_268delinsCC [p.Ala88Profs*51]; c.55dupG [p.Glu19Glyfs*14]) leading to absence of protein expression (HMOX1 ^MUT/MUT^)(19) and two healthy controls (HMOX1 ^WT/WT^). Samples were mixed with washing medium (RPMI 1640 (# 21875034) containing 1 × antibiotic-antimycoticum (#15240062)). Cells were isolated by centrifugation at 500 × g for 10 min followed by incubation with the supernatant of the EBV-producing marmoset cell line B95-8 (49) in order to establish LCL. Cells were incubated with culture medium consisting of RPMI 1640, 1 x antibiotic-antimycotic and 20% (vol/vol) fetal bovine serum (#10270106) at 37 °C and 5% CO_2_.

As a variant-independent model, a CRISPR/Cas9-mediated knock out of *HMOX1* in HEK293T cells was performed as previously described (50) (HEK293^HO-1-/-^). Wild-type HEK293T cells were purchased from Abcam (#ab255449, Abcam, Cambridge, United Kingdom). HEK cells were maintained in DMEM + D-Glucose + L-Glutamine + Pyruvate (#11995065) containing 10% (vol/vol) FBS, and 1 x antibiotic-antimycotic at 37 °C in an atmosphere containing 5% CO_2_.

All cells were genotyped before performance of experiments and regularly tested for mycoplasma contamination using isothermal PCR with the MycoStrip – Mycoplasma Detection Kit (#rep-mys-20, InvivoGen, San Diego, CA, USA) according to the manufacturer’s protocol.

### Flow cytometry

Cells were washed with DPBS and Annexin V Binding Buffer (#422201 BioLegend, San Diego, CA, USA) followed by staining with 400 µl of Annexin V Binding Buffer, 4 µl of APC Annexin V (#640941, BioLegend) and 4 µl of 7-AAD (#420404 BioLegend). Cells were incubated in the dark on ice for 15 min. Flow cytometry was performed using a BD FACSCanto flow cytometer (BD Scientific, Franklin Lakes, NJ, USA). A total of 10,000 events per sample were acquired and displayed in BD FACSDiva software. Compensation for both antibodies was performed on single stained cells.

To determine cell viability, we calculated the percentage of viable cells relative to the total number of cells analyzed. This was achieved by gating viable cells based on their forward scatter and negative staining for both annexin V and 7-AAD. FlowJo (version 10.8.0) was used for data analysis.

### Hemin cytotoxicity assay

To model the effect of a loss of HO-1 function in the presence of the enzyme substrate, patient-derived and wild-type lymphoblastoid cells (2.5 × 10^5^ cells/ml) were exposed to ascending concentrations (0 µM, 50 µM, 75 µM, 100 µM) of porcine hemin (# 51280) for 24 h under standard conditions (10). Wild-type and knock-out HEK293T cells were incubated with 0 µM, 15 µM, 30 µM, and 60 µM of hemin.

Viability was determined by flow cytometry as stated above.

### Transmission electron microscopy of HO-1 deficient cell lines

For ultra-structural analysis of mitochondria, 2.5 x 10^5^ cells of LCL and HEK293T respectively were incubated for 24 h either with or without 25 µM hemin before washing with PBS and mixing in 1 ml glutaraldehyde solution (2.5% v/v in Sörensen phosphate buffer). Samples were fixed with 1% OsO_4_, dehydrated, and embedded in glycid ether 100. 60 nm ultrathin sections were cut using a Leica Ultracut R ultramicrotome (Leica Microsystems, Wetzlar, Germany) and counterstained with 8% uranyl acetate in bidistilled water and incubated with lead citrate solution. Samples were inspected with a CM 10 transmission electron microscope (Philips, Amsterdam, Netherlands). Mitochondrial morphology assessment in wild-type and HO-1 deficient LCL, as well as wild-type and HO-1 knockout HEK293T cells was performed following a standardized protocol for TEM image analysis (51) with Fiji (52). Cells were imaged and analyzed by separate, blinded individuals (V.V.M. and B.H. - imaging, P.S.W. - measurement).

### Respiratory chain complex activity measurements

The activity of the individual respiratory chain complexes was assessed in HEK293 ^HO-1^ ^+/+^ and HEK293 ^HO-1-/-^ cells cultured under standard conditions as outlined above. For spectrophotometric analysis of respiratory chain complexes, cells were washed twice with HEPES buffer (NaCl 110 mM, KCl 2.6 mM, KH_2_PO_4_ 1.2 mM, CaCl_2_ 1 mM, MgSO_4_ 1.2 mM, HEPES 25 mM, pH 7.4), followed by incubation in this buffer supplemented with 10 mM glucose for 15 min at room temperature. Following sonication for 2 × 10 s (single pulses of 0.3 s at 0.7s intervals, 20 W), respiratory chain enzyme activities of complexes I–V were determined spectrophotometrically as previously described (53).

### Cellular energetics measurements

To assess the oxygen consumption rate (OCR), an indicator of mitochondrial respiration, and the extracellular acidification rate (ECAR), an indicator of aerobic glycolysis, the Seahorse XFe 96 analyzer (#101991-100, Agilent, Santa Clara, CA, USA) was used according to the manufacturer’s recommendations. Briefly, HEK293 ^HO-1^ ^+/+^ and HEK293 ^HO-1-/-^ were seeded at a density of 2.5 × 10^4^ cells/well on a poly-D-lysine (#P6407-5MG) coated Seahorse XF96/XF Pro cell culture microplate (#204624-100, Agilent) on the day prior to the analysis. Cells were preincubated in a CO2 free incubator at 37 °C for 1 h in Seahorse XF DMEM medium, pH 7.4 (#103575-100, Agilent) substituted with 10 mM glucose (#G5767-500G), 2 mM glutamine (#G3126-100G), and 1 mM pyruvate (#P2256-25G).

The mitochondrial stress and glycolytic rate test were performed at final inhibitor concentrations of 2.0 µM oligomycin (#O4876-5MG), 1 µM carbonyl cyanide-p-trifluoromethoxyphenylhydrazone (FCCP, #C2920), 0.1 µM rotenone (#R8875), 1 µM antimycin A (#A8674), and 50 mM 2-deoxy-D-glucose (2-DG, #D8375-25G). Following baseline measurements, three measurements per injection were taken. Results were normalized to cell counts as determined by Hoechst 3342, trihydrochloride, trihydrate (#H3570, Thermo Fisher) staining at a concentration of 5 µg/ml followed by fluorescence measurement using a Tecan Infinite 200 PRO plate reader (Tecan Group AG, Männedorf, Switzerland) with 355/9 nm excitation and 460/20 nm emission filters.

### Quantification of cellular ATP

Cellular ATP content was assessed using the CellTiter-Glo luminescent assay (#G7570, Promega, Madison, WI, USA) according to the manufacturer’s protocol. Briefly, 5000 cells/well were seeded in DMEM in a 96-well plate. Simultaneously, an ATP standard curve was generated with ascending concentrations (0.05, 0.1, 0.25, 0.5, 1, and 2.5 µM) of ATP (#K054.1, Carl Roth GmbH + Co. KG, Karlsruhe, Germany). 5050 µl of CellTiter-Glo reagent were added after incubation at 37 °C in an atmosphere containing 5% CO_2_ overnight, followed by luminescence measurement using a TriStar2 LB 942 microplate reader (Berthold Technologies GmbH & Co. KG, Bad Wildbad, Germany) with a counting time of 0.1s.

Luminescence was normalized to cell number as determined by Hoechst 33342 staining.

### Evaluation of antioxidant compounds

To study the effects of various antioxidants on cell survival following hemin stress, cells were seeded at a density of 2.5 × 10^5^ cells/ml and treated with antioxidants at concentrations typically reached with standard doses or solute alone (Tab. 2). After a pre-incubation period of 1 h, 75 µM hemin were added and cells were incubated for another 24 h followed by flow cytometric analysis.

The effect of 2-Phospho-L-ascorbic acid trisodium salt (AA2P) on the viability of hemin-stressed LCL ^HO-1MUT/MUT^ was assessed in a dose escalation study. Cells were treated with concentrations of AA2P ranging between 70 µM and 49 mM and incubated for one hour followed by 24-hour incubation with 75 µM hemin. The minimum and maximum concentrations of AA2P were chosen based on data originating from previous pharmacokinetic studies. A plasma concentration of 70 µM Vitamin C can be achieved from a healthy standard diet (54) and results in the sodium-dependent vitamin C transporter 2 (SVCT2) reaching its V_max_ (23, 55, 56). The maximum plasma concentration of 49 mM was achieved by an intravenous administration of 1 g/min Vitamin C at four consecutive days for four weeks (57).

### Inhibition of SVCT2

To distinguish between intra- and extracellular effects of AA2P, uptake via SVCT2 was inhibited using sulfinpyrazone (#S9509) (20). To this end, LCL were incubated with 500 µM of sulfinpyrazone for 10 min, followed by 1 h incubation with 1 mM AA2P under standard cell culture conditions. 75 µM hemin was added to the cells and after another 24 h, viability was assessed via flow cytometry.

### LC-MS/MS of antioxidant metabolites

Three replicates of dry cell pellets and cleared conditioned culture media for each experimental group were isolated and stored at −80 °C until the day of sample preparation for metabolomic measurement. Whole cell lysates were prepared as described previously (43). Sulfur-containing metabolites as well as creatinine were determined according to a previously published procedure (43, 58). Calibration curves for all metabolites were prepared from individual stock solutions prepared in house. Quantitation accuracy was examined by monitoring homocysteine and methylmalonic acid concentrations in an external quality control, namely, the Control Special Assays in Serum, European Research Network for the evaluation and improvement of screening, diagnosis, and treatment of Inherited disorders of Metabolism (ERNDIM) IQCS, SAS-02.1 and SAS-02.2 from MCA Laboratories, Winterswijk, Netherlands. For all other metabolites, quantitation trueness was tested by examining metabolite concentrations in plasma from a previously validated sample isolated from a healthy control individual with respect to standard reference ranges, using the same calibration curves and LC-MS/MS running conditions.

Metabolite concentrations were normalized by total protein concentration of the whole cell lysate, and unless otherwise indicated, expressed as nmol metabolite/mg protein. Total protein concentration was determined using the bicinchoninic acid assay (#23225, Thermo Fisher). Signal processing and quantification of metabolites was carried using Analyst (version 1.7.2, AB Sciex, Darmstadt, Germany).

### Immunoblotting

For whole cell lysates, 1 × 10^7^ cells were washed with ice-cold PBS, followed by incubation with RIPA buffer containing 150 mM NaCl, 1.0% IGEPAL CA-630, 0.5% sodium desoxycholate, 0.1% SDS, 50 mM Tris, pH 8.0) for 30 min on ice. Adhesive cells were scraped after addition of the buffer. The suspension was centrifuged at 10 000 × g for 10 min at 4 °C. The supernatant was diluted to a concentration of 1 µg protein / µl in RIPA buffer and mixed with LI-COR 4 × protein sample loading buffer (#928-40004, LI-COR Biosciences, Lincoln, NE, USA), followed by denaturation at 95 °C for 5 min. For SVCT2 detection, 10 µg of protein were separated on 10% Mini-PROTEAN TGX Stain-Free Protein Gels (#4568036, Bio-Rad) according to the manufacturer’s instructions. Total protein visualization and transfer onto PVDF membranes was carried out following the manufacturer’s protocol. Membranes were blocked with 5% milk in TBST for 1 h at room temperature and constant agitation, followed by incubation with primary antibodies in blocking buffer at 4 °C overnight.

SVCT2 Polyclonal Antibody (#27019-1-AP, Proteintech, Rosemont, IL, USA) was used at a dilution of 1:5000 in 5% milk in TBST. HRP-conjugated Affinipure Goat Anti-Rabbit IgG(H+L) (#SA00001-2, Proteintech, dilution 1:5000) was used as a secondary antibody. Chemiluminescent visualization was carried out using the Cytiva Amersham ECL Prime Western Blotting Detection Reagent (#10308449, Cytiva, Marlborough, MA, USA) as per the manufacturer’s instructions. Blots were visualized on a Chemidoc apparatus. Six biological replicates per genotype were analyzed.

Densitometric analyses were performed using Image Lab software (Bio-Rad) with normalization to total protein as determined by stain-free technology.

### Statistics and software

All statistical analyses were performed using GraphPad Prism (v10.2.3, GraphPad Software, Inc., La Jolla, CA, USA). Means of two groups were compared using the two-sided nonparametric Mann-Whitney *U* test or student’s unpaired two-sided t-test, depending on the distribution of data. One- or two-way analysis of variance (ANOVA) were used for comparison of more than two groups, followed by Dunnett’s and Šidák’s post-hoc test respectively. Multiple non-normally distributed groups were compared using the Kruskal-Wallis test followed by Dunn’s post-hoc test. Data are presented as mean ± s.e.m. and *P* values <0.05 were considered to be statistically significant. For normalization of data, values were divided by the mean of the respective control group, i.e. either solute-incubated or wild-type cells. Relative viabilities were calculated by dividing through the mean of the respective control group and dividing by 100. If not indicated otherwise, experiments were conducted with *n* = 3 biological as well as three technical replicates. Figure 3 as well as the graphical abstract were created with biorender.com.

### Study approval

Written informed consent of the participants or their legal guardians was obtained prior to inclusion in the study. The study was approved by the relevant IRB (Ethikkommission der Ärztekammer Westfalen-Lippe und der Westfälischen Wilhelms-Universität Münster, reference number: 2021-289-f-S) and was performed in line with the principles of the Declaration of Helsinki.

## Data availability

Source data for the graphs are provided in the Supporting Data Values file. Unedited blots are available in the Supplemental Material. No original code was generated in this study. Any additional information required to reanalyze the data reported in this paper is available from the lead contact upon request.

## AUTHOR CONTRIBUTIONS

L.-S.B.: investigation, methodology, validation, data curation, writing - original draft, writing - review & editing; P.S.W.: investigation, methodology, validation, writing - review & editing; A.M.T.: investigation, methodology, writing - review & editing; V.W.: investigation, data curation, writing - review & editing; S.B.: investigation, writing - review & editing; A.S.: investigation, methodology, writing - review & editing; V.V.M.: investigation, methodology, writing - review & editing; B.H.: investigation, methodology, writing - review & editing; H.W.: investigation, data curation, writing - review & editing; A.M.D.: investigation, methodology, writing - review & editing; L.H.: investigation, methodology, validation, formal analysis, writing - review & editing; W.N.N.: methodology, data curation, writing - review & editing; A.J.: methodology, data curation, writing - review & editing; L.K.: conceptualization, methodology, writing - review & editing; G.V.: methodology, validation, data curation, writing - review & editing; T.M.: investigation, writing - review & editing, conceptualization; J.H.P.: investigation, methodology, validation, data curation, formalization, visualization, writing - original draft, writing - review & editing, conceptualization, supervision.

## Supporting information

Unedited blots

Supporting Data Values

## ACKNOWLEDGEMENTS

We are indebted to the patient’s family for their support and dedication throughout the conception of the manuscript. The expert technical assistance of Susanne Schleifenbaum, Melanie Saers, Andreas Borgscheiper, Martina Herting, and Kai Wohlgemuth as well as Cordula Westermann and Astrid Fitter is gratefully acknowledged. We thank Birgit Burkhard, Thomas Kaiser, and Heymut Omran as well as Akihiro Yachie for fruitful discussions during the treatment of the patient and conception of this work. JHP is supported by a non-restricted stipend from the North Rhine-Westphalian Academy of Arts and Sciences (Nordrhein-Westfälische Akademie der Wissenschaften und der Künste) as well an Else Kröner Memorial Fellowship (2022_EKMS.06) by the Else Kröner-Fresenius-Stiftung (EKFS). This research was funded by a grant by Innovative Medical Research (IMF), Medical Faculty University of Münster (PA 1 2 23 08). Several authors of this publication are members of the European Reference Network for Rare Hereditary Metabolic Disorders (MetabERN).

## Conflict-of-interest statement

The authors declare that no conflict of interest exists.

---

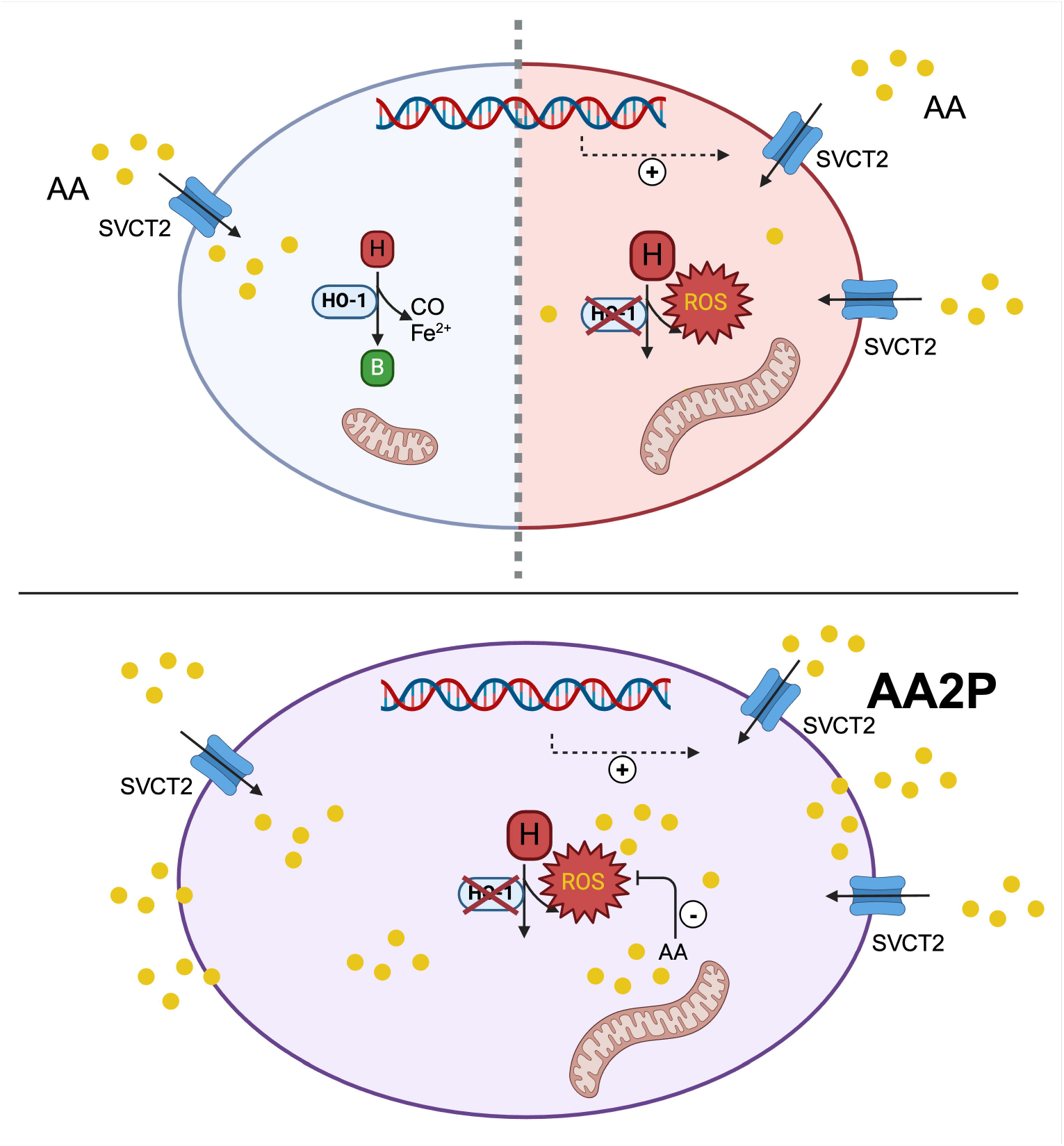

## Notes

### Competing Interest Statement

The authors have declared no competing interest.

