## Supplemental for "Treatment with 2-phospho-L-ascorbic acid mitigates biochemical phenotypes of heme oxygenase 1 deficiency"

#### SUPPLEMENTAL METHODS

##### RNA sequencing

Triplicate cultures of LCL $^{\text{HO-1 MUT/MUT}}$ as well as in LCL $^{\text{HO-1 WT/WT}}$ were incubated with and without AA2P at a concentration of 1 mM followed by mechanical dissociation and centrifugation of cells at 500 × g and 4 °C. Pellets were snap frozen in liquid nitrogen and stored at -80 °C until further processing.

RNA was isolated using the QIAGEN RNeasy Mini kit (#74104, QIAGEN, Hilden, Germany) according to the manufacturer's protocol. Following RNA quality control by a fluorescence based quantification method and fragment length analysis, library preparation was performed using the TruSeq Stranded mRNA kit (#20020594, Illumina, San Diego, CA, USA). After sequencing on a NovaSeq 6000 system (Illumina, sequencing parameters: 2 × 100 bp), demultiplexing of the sequencing reads was performed with Illumina bcl2fastq (2.20). Adapters were trimmed with Skewer (version 0.2.2) (*1*). Trimmed raw reads were aligned to hg19-cegat using STAR (version 2.7.3) (*2*). Differential expression analysis between groups was performed with DESeq2 (version 1.24.0) (*3*) in R (version 3.6.1). DESeq2 uses a negative binomial generalized linear model to test for differential expression based on gene counts. The raw counts derived from the mapping contain the number of reads that map to each geneID. Based on these numbers the normalized counts are calculated. In the first step of the normalization, DESeq2 was used to calculate a fictive "reference sample" defined as the geometric mean for each gene across all samples regardless of group affiliation. The counts for each gene and sample were then divided by this reference value.

In the next step the size factor was estimated for each sample by calculating the median of these ratios. To get the normalized counts, for each gene and sample the raw counts were divided by the sample's size factor. This normalization accounts for different library sizes and for biases if, e.g.,in some samples only a few genes are very highly expressed. As recommended by DESeq2, genes with less than two reads over all samples were removed. Using normalized counts, DESeq2 was used to calculate the log2 fold change, where the p-value reports the statistical significance of this result (Wald test). The Benjamini-Hochberg correction was used to correct for multiple testing, generating adjusted *P* values (padj).

The quality of the FASTQ files was analyzed with FastQC (version 0.11.5-cegat) (*4*). Plots were created using ggplot2 (*5*) and ggdendro (version 0.1.23) (*6*) in R (version 3.6.1).

The heatmap shows the 50 most differentially expressed genes (25 upregulated, 25 downregulated). The z-score of normalized counts was used to create the heatmap. The columns and lines are arranged in such a way that genes and samples with similar expression cluster together. Genes with log2 Fold change > |1.5| and adjusted p-value < 0.05 are marked as significantly differentially expressed.

### SUPPLEMENTAL FIGURES

#### Supplemental Figure S1

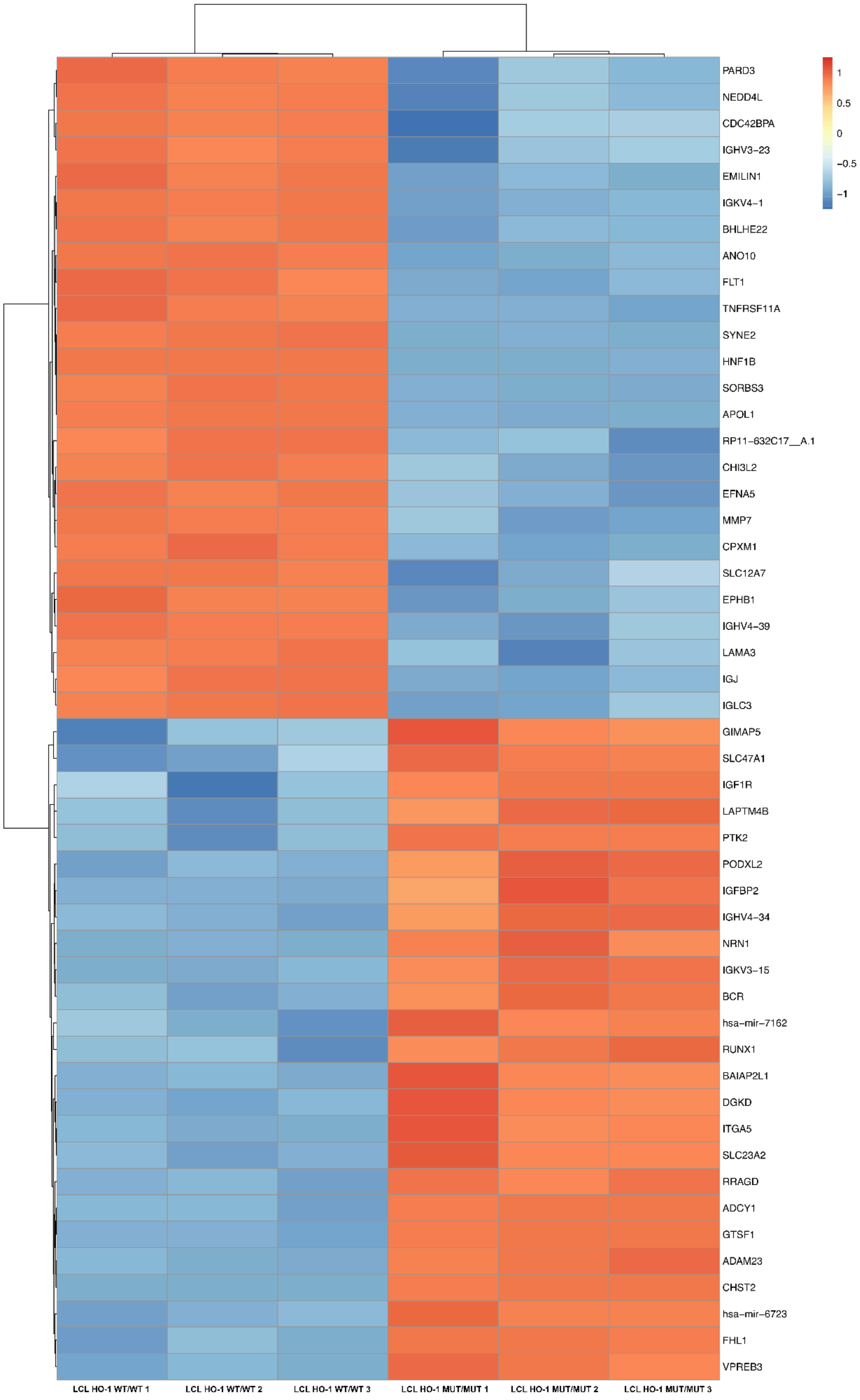

**Supplemental Figure S1 - Identification of differentially expressed genes in HO-1 deficiency**
Differential gene expression analysis using RNA Sequencing was performed in LCL $^{\text{HO-1 MUT/MUT}}$ in comparison to LCL $^{\text{HO-1 WT/WT}}$. The heatmap shows the 50 most differentially expressed genes (25 upregulated, 25 downregulated). The z-score of normalized counts was used to create the heatmap. The columns and lines are arranged in such a way that genes and samples with similar expression clustered together. Genes with log2 fold change > |1.5| and adjusted p-value < 0.05 are marked as significantly differentially expressed. N=3 replicates per genotype were analyzed.

#### Supplemental Figure S2

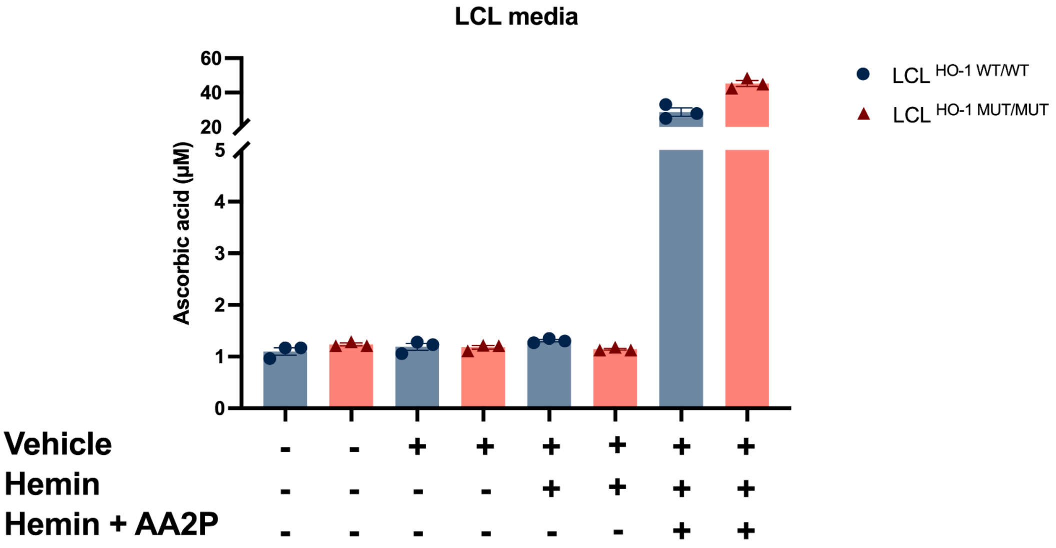


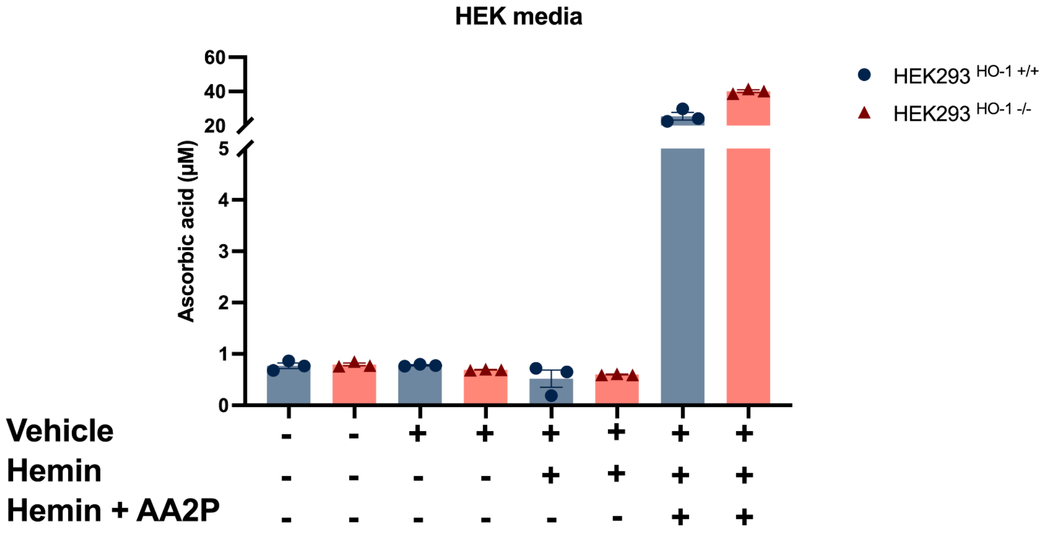


**Supplemental Figure S2 - Ascorbic acid content of cell culture media**
LC-MS/MS was used to quantify the ascorbic acid concentration in cell culture media of cells in which intracellular ascorbic acid concentrations were analyzed (Fig. 2 + 5 of the main manuscript. Ascorbic acid concentrations were not different between wild-type and HO-1 deficient LCL and HEK293T cells and remained stable upon addition of 75 µM hemin to the culture media. Addition of 1 mM 2-phospho-L-ascorbic acid led to a steep increase in ascorbic acid concentration with no statistically significant difference between wild-type and HO-1 deficient cells. Data are mean ± s.e.m. Means were compared between genotypes using the Kruskal-Wallis test followed by Dunn's post-hoc test. *N* = 3 replicates.

4. S. Andrews, *FastQC: A Quality Control Analysis Tool for High Throughput Sequencing Data* (Github; https://github.com/s-andrews/FastQC).

5. H. Wickham, *ggplot2* (Springer New York; https://link.springer.com/book/10.1007/978-0-387-98141-3).

6. A. de Vries, *NEWS.md at Main · Andrie/ggdendro* (Github; https://github.com/andrie/ggdendro/blob/main/NEWS.md).
