## Supplementary material for "Treatment with 2-phospho-L-ascorbic acid mitigates biochemical phenotypes of heme oxygenase 1 deficiency": Unedited blots

Full unedited blots

Berendes et al.

Figure 2

LCL - Total protein stain

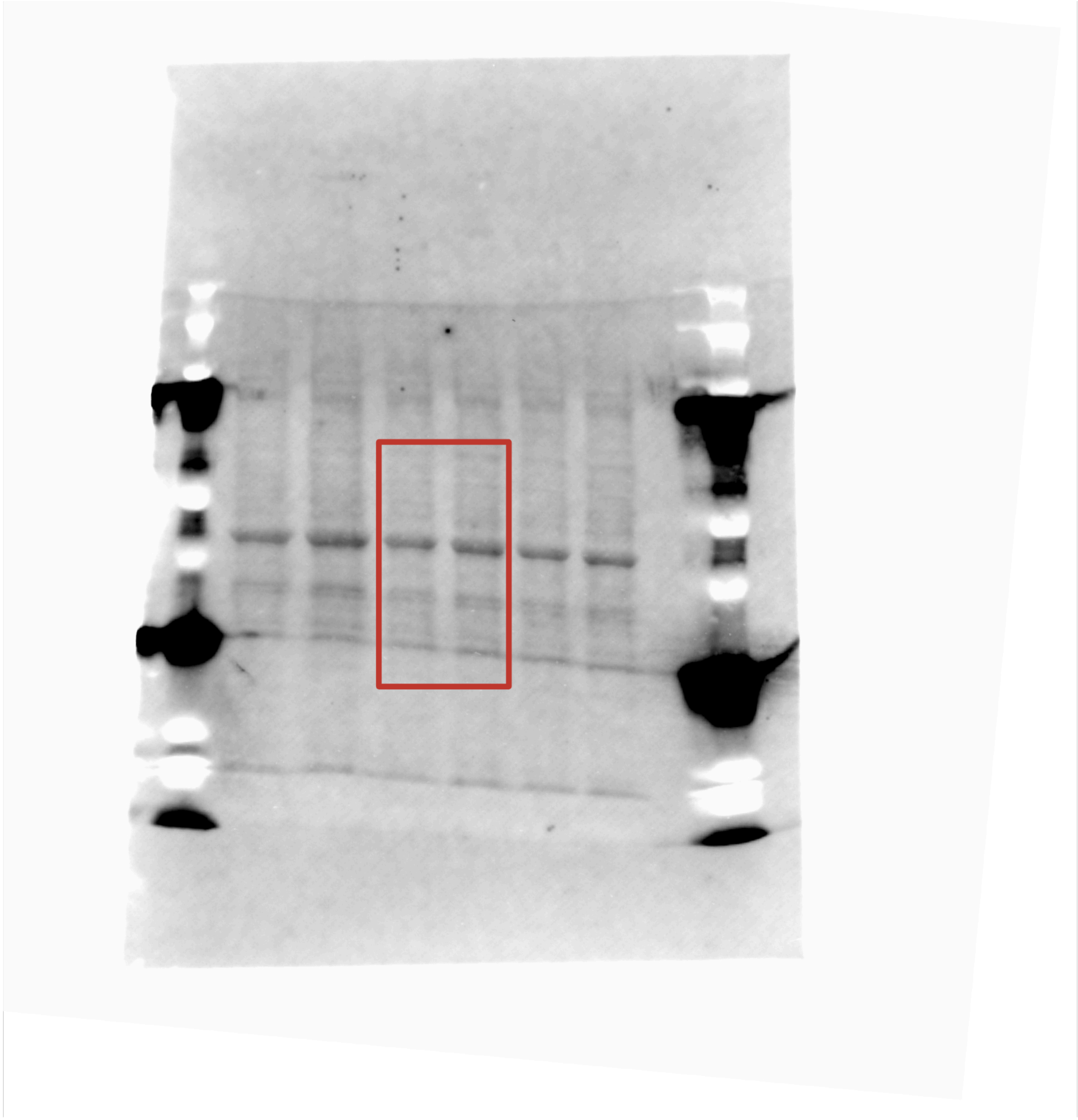

LCL - SVCT2

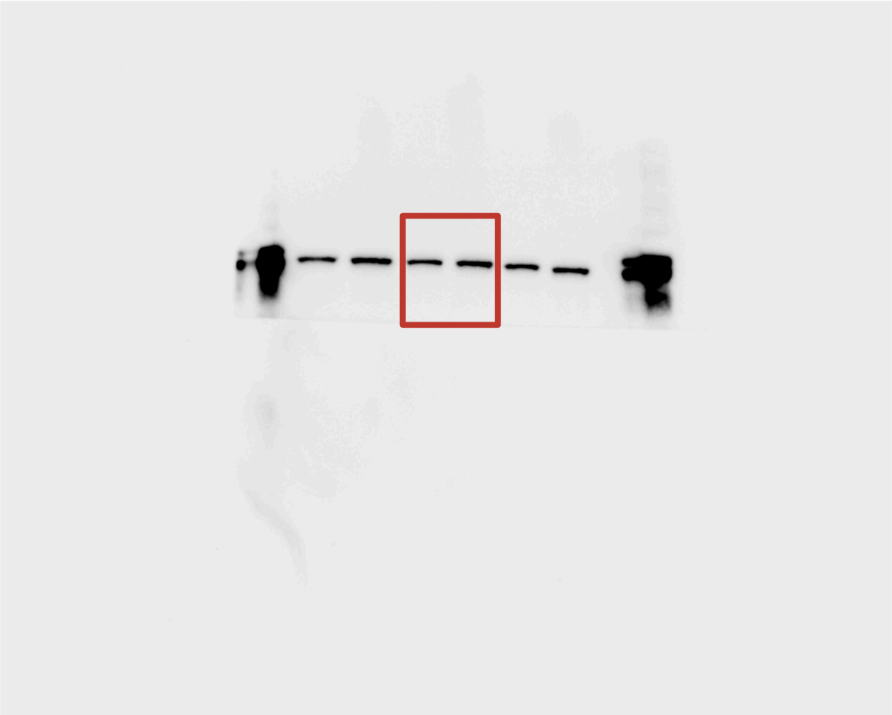

HEK293T - Total protein stain

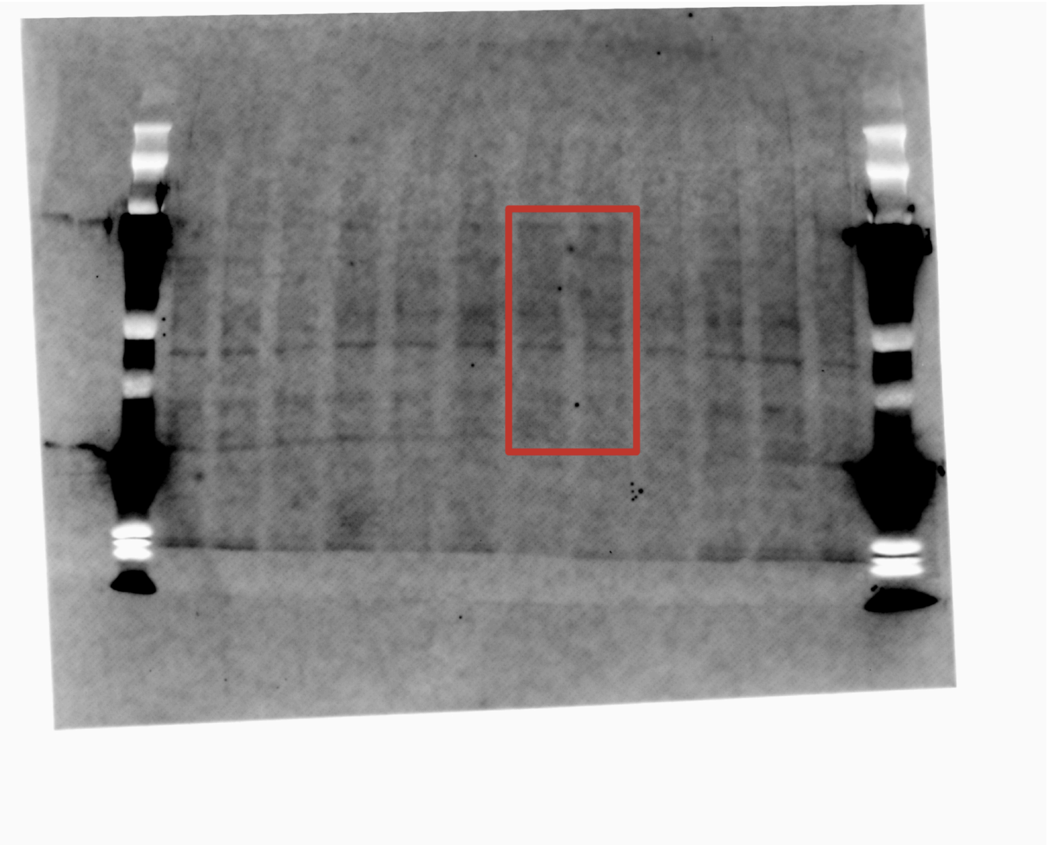

HEK293T - SVCT2

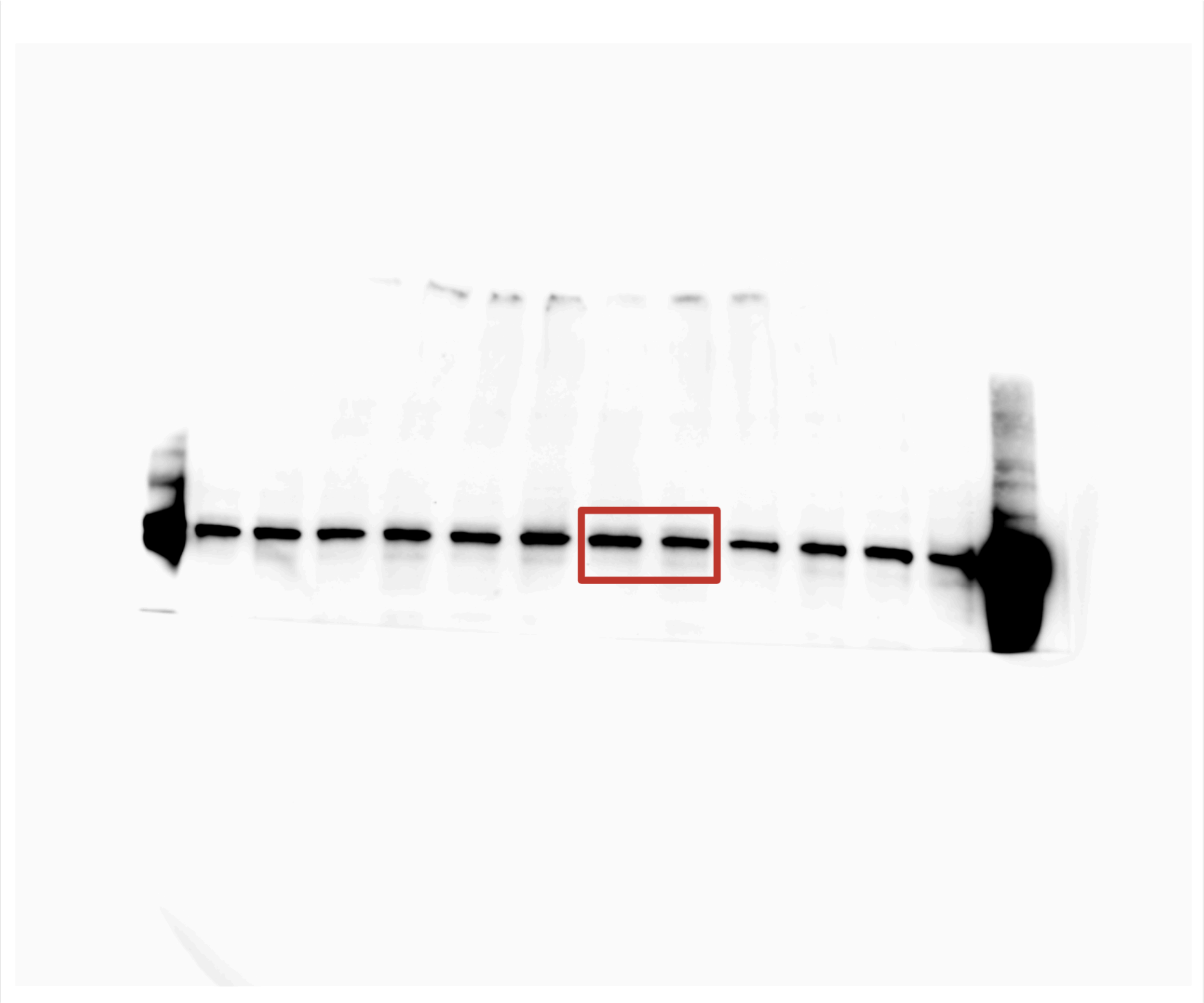
